# microRNA regulation of persistent stress-enhanced memory

**DOI:** 10.1101/379594

**Authors:** Stephanie E. Sillivan, Sarah Jamieson, Laurence de Nijs, Meghan Jones, Clara Snijders, Torsten Klengel, Nadine F. Joseph, Julian Krauskopf, Jos Kleinjans, Christiaan H. Vinkers, Marco P.M. Boks, Elbert Geuze, Eric Vermetten, Kerry J. Ressler, Bart P.F. Rutten, Gavin Rumbaugh, Courtney A. Miller

## Abstract

Disruption of persistent, stress-associated memories is relevant for treating posttraumatic stress disorder (PTSD) and related syndromes, which develop in a subset of individuals following a traumatic event. Using a stress-enhanced fear learning protocol that results in differential susceptibility in inbred mice, we integrated small-RNA sequencing with quantitative proteomics on basolateral amygdala tissue collected one month after training. We identified persistently changed microRNAs, including mir-135b-5p, and predicted target proteins associated with PTSD-like heightened fear expression. Functional manipulations of mir-135b-5p bidirectionally modulated stress-associated memory. mir-135b-5p is expressed in human amygdala and its passenger strand was elevated in serum from a well-characterized military PTSD cohort. miR-135b-5p is a therapeutic target for dampening persistent, stress-enhanced memory and its passenger strand a potential biomarker for responsivity to a mir-135-based therapeutic.

**One Sentence Summary:** mir-135 can be manipulated to weaken persistent, stress-associated memory and serve as a biomarker of PTSD.

## Main Text

Unlike memory acquisition and consolidation, mechanisms supporting long-lasting, remote memory are largely unknown, yet highly relevant to neuropsychiatric disorders of memory, such as posttraumatic stress disorder (PTSD). PTSD is a chronic, debilitating disorder in which patients exhibit memories of trauma that are heightened, perseverant and extinction-resistant (*1*). Nearly everyone experiences at least one traumatic event in their lifetime, but only 10-20% will later display enduring symptoms of PTSD (*2*). Exposure therapy, a form of cognitive behavioral therapy (CBT) that utilizes extinction of fearful memories, is considered the gold standard PTSD treatment. However, some patients are resistant or experience exacerbated symptoms and, of those who do respond, most retain their PTSD diagnosis (*3*). Development of adjunctive pharmacotherapies to enhance success of CBT is needed, but deeper insight into the neurobiology of PTSD-like behaviors to identify potential targets is first required. To identify mechanisms governing differential stress susceptibility, we developed a stress-enhanced fear learning (SEFL) paradigm in inbred C57BL/6 mice that results in PTSD-like characteristics, including persistently enhanced memory in a subset of mice termed stress-susceptible (SS) (*4*). This contrasts with stress-resilient (SR) mice that display no stress-induced enhancements. Importantly, this SEFL protocol allows for the study of molecular phenotypes associated with selective vulnerability, as SS mice can be identified from training data, avoiding mechanistic confounds introduced by additional phenotyping. We also reported differential expression of genes with known polymorphisms in human PTSD genomic studies between SS and SR animals (*4*).

A role for miRNAs in mediating synaptic plasticity and learning has emerged over the past decade (*5*). miRNAs are endogenous ~20-24 nucleotide RNAs that act as translational repressors through direct binding to the 3’-UTR of target mRNAs and noncleavage degradation of target mRNA via deadenylation (*6*). Protein translation is required for formation of new memories, and its regulation via synapse-specific miRNA localization can modulate synaptic plasticity (*7–9*). Indeed, expression of miRNA biogenesis regulators Dicer and Pasha (Dgcr8) impacts fear memories (*10, 11*). Both cued and contextual fear conditioning (FC) regulate miRNA expression and modulation of such miRNAs impacts consolidation and retrieval (*12–17*). These prior studies assayed recent fear memory with no stress component. The contribution of miRNAs to long-lasting, stress-enhanced remote (>30 days) memory is unknown. miRNAs are excellent remote memory candidate regulators because their wide genomic range of target proteins confers a complexity capable of handling the intricacies of memory. Importantly, mature miRNAs can have very long half-lives (i.e. months) (*18*), lending themselves to the goal of memory persistence.

Here we examined involvement of miRNAs in mediating susceptibility to long-lasting traumatic memories in our SEFL model, with the hypothesis that a unique miRNA signature defines stress-enhanced memory and differentiates it from stress resilience. SEFL was performed in male mice, followed by a remote memory test 30 days later (**Fig. 1A**). As we have previously reported (*4*), prior stress resulted in varied memory strength at recall that was predicted by freezing during FC (**Fig. S1A**). Indeed, stressed animals that froze more during training displayed the highest freezing during the remote memory recall test and were classified as SS. To interrogate mechanisms underlying differential memory strength between SS and SR animals, despite identical training and genetic homozygosity, we performed small RNA-sequencing (smRNA-Seq) and quantitative proteomics on basolateral amygdala complex (BLC; lateral and basolateral) tissue of SEFL-trained mice. Tissue was isolated 30 days after training (no retrieval test) to identify miRNAs in SS mice contributing to enhanced memory strength through persistent change following SEFL, not miRNA changes dynamically induced by the act of memory retrieval. The BLC was chosen because long-term fear memories rely on this region (*19*) and we previously demonstrated that SS animals have exaggerated Fos activation in the BLC relative to SR and FC (no stress) animals, consistent with human PTSD studies (*20*). Importantly, freezing at the end of training, which classifies stress susceptibility, did not differ between SR and FC animals, but did differ between SR and SS (**Fig. S1B**).

**Fig. 1.**
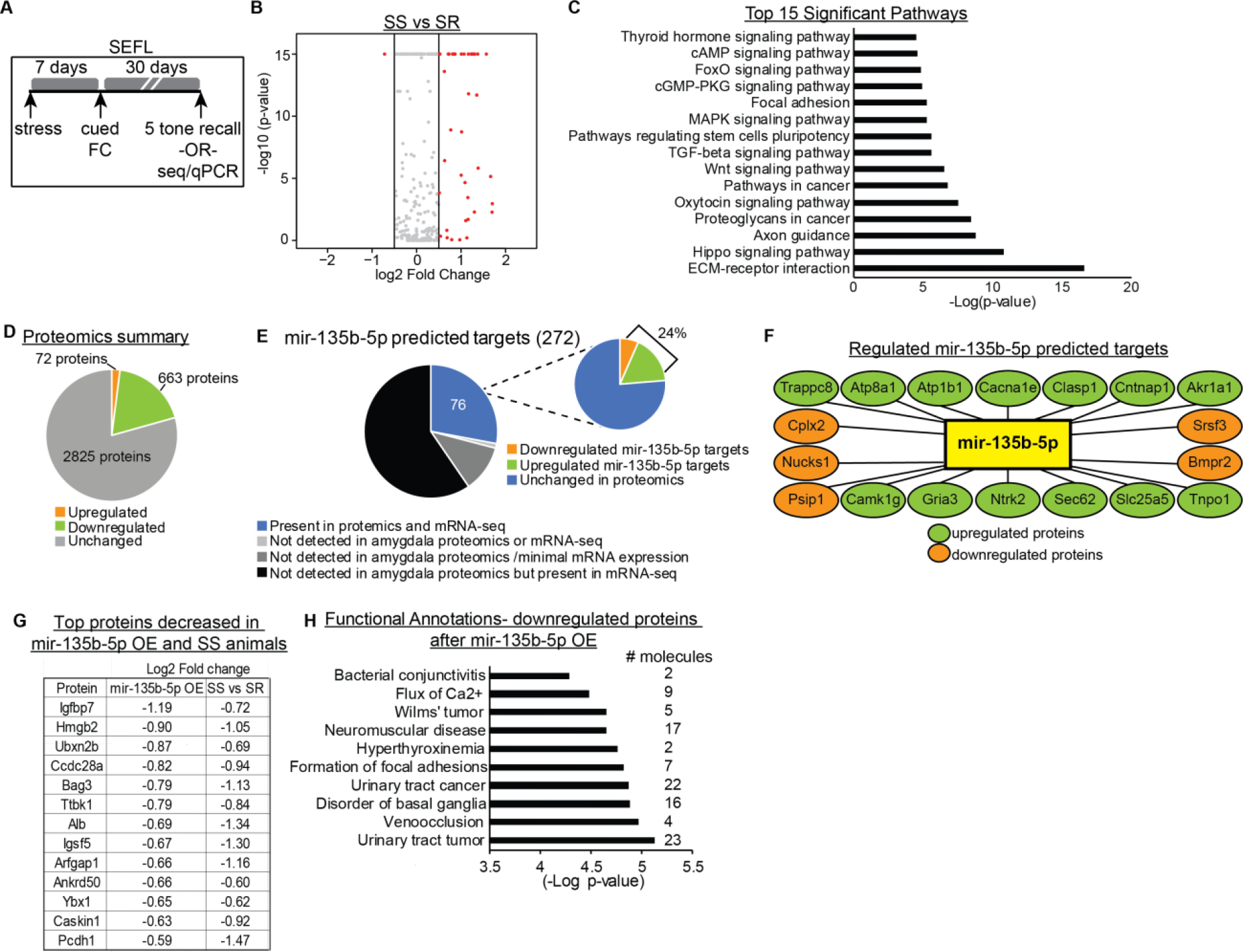
Long-lasting miRNA and protein changes in the BLC following SEFL in stress susceptible mice. (A) Overview of SEFL paradigm. (B) Volcano plot of miRNA expression in SS vs SR animals. (C) DIANA mirPath analysis to identify pathways significantly targeted by SS miRNAs. (D) Summary of protein changes in BLC tissue between SS and SR animals 30 days after SEFL. (E-F) Proteomics expression of putative mir-135b-5p targets between SS and SR animals. (G-H) Proteins changed by >50% in both SEFL and premirOE-135b-5p-naïve animals and top regulated pathways of proteins changed by >25% in those groups.

SmRNA-Seq was performed on two separate cohorts of animals at two genomics facilities employing different reagents and bioinformatics pipelines to yield miRNA profiles associated with susceptibility to stress-enhanced memory. We identified 42 miRNAs differentially expressed between SS and SR groups in both datasets, of which 41 were upregulated in SS animals (**Fig. 1B, Table S1 and SE.1-2**). Pathway analysis with DIANA’s mirPATH software using the microCts algorithm identified target pathways predicted to be regulated by these differentially expressed miRNAs. It included pathways that critically regulate stress and learning processes, including ECM-receptor interaction, MAPK signaling pathway and Thyroid Hormone Signaling (**Fig. 1C**). DIANA mirPath also provides access to two other target predicting algorithms, TargetScan and Tarbase. We compared all three algorithms and used the results to guide sequencing validation to miRNAs predicted by at least 2 algorithms that have targets significantly involved in pathways related to learning and memory (**Table S2**). One candidate, mir-135b-5p, is predicted to target proteins in both ECM-receptor interaction and Thyroid Hormone Signaling pathways. Interestingly, thyroid hormone signaling has been implicated in amygdala-based memory processes (*21, 22*) and family members of mir-135b-5p have been studied in depressive and anxiety-like behaviors (*23, 24*), cortical axon guidance (*25*) and synaptic plasticity (*26*). Importantly, mir-135b-5p is conserved from mouse to human brain tissue (*27*). Only one study has examined mir-135b-5p in the context of neuronal processes, identifying both mir-135a and mir-135b as regulators of axon guidance, migration and regeneration in cultured cortical and hippocampal cells (*25*).

miRNAs are appealing to study in the context of psychiatric disorders because each one has potentially hundreds of protein targets. Therefore, we performed quantitative mass spectrometry (MS) on SEFL animals 30 days after training (no extinction) to identify potential mir-135b-5p targets that may participate in stress-enhanced memory. 3,560 proteins were detected by MS, with 72 downregulated and 663 upregulated by at >1.5 log2 fold change between SS and SR groups (**Fig. 1D and SE.3**). The top 10 most significant pathways from Ingenuity Pathway Analysis (IPA) revealed functional annotations related to neuronal structure (e.g. Density of microtubules, Organization of cytoskeleton) in the downregulated protein list (**Fig. S2A**), while the upregulated pathways were primarily related to cancer (e.g. Adenocarcinoma, Digestive system cancer) (**Fig. S2B**). However, the 5 proteins with the greatest increase in SS BLC included proteins with known roles outside the cancer field (**Fig. S2C**). While many proteins are expected to participate in miRNA-independent processes, downregulation of SS proteins could reflect a coordinated interaction with upregulated miRNAs responsive to stress-enhanced memory, such as mir-135b-5p. A total of 272 mir-135b-5p targets were predicted by at least two of the three major databases (TargetScan, DIANA, mirDB), but only 76 were detected in BLC tissue at both the RNA and protein level (**Fig. 1E**). Interestingly, 24% (*18*) of predicted mir-135b-5p protein targets detected in the BLC were regulated between SS and SR animals and included Ntrk2, a BDNF receptor well-characterized for its role in amygdala-dependent learning and PTSD (**Fig. 1E-F**) (*28*). Pathway analysis of these 18 mir-135b-5p putative targets that were changed by SEFL identified annotations related to neuronal function, including spatial learning, long-term depression of neurons, plasticity of synapse and excitatory postsynaptic potential of neurons (**Fig. S2D**). Because many predicted mir-135b-5p targets could be false positives, we also examined the BLC proteome of naïve animals injected with a lentivirus to overexpress (OE) mir-135b-5p in the BLC at physiologically relevant levels (premirOE-135b-5p; **Fig. S3A-B**). We aligned the MS datasets and identified 13 proteins downregulated by >50% in both SS animals and naive premirOE-135b-5p animals (**Fig. 1G**), finding 99 proteins downregulated by >25%. IPA of these downregulated proteins identified pathways related to neuronal function (e.g. Neuromuscular disease, Disorder of basal ganglia), as well as non-neuronal pathways with molecules in need of functional studies in the brain (**Fig. 1H**).

We examined mir-135b-5p in the adult mouse brain and found it is broadly expressed (**Fig. 2A**). In situ hybridization revealed anatomical localization of mir-135b-5p throughout the BLC (**Fig. 2B**). Using a cell fractionation assay, mir-135b-5p expression was detected in synaptosomes, where it could regulate synaptic protein translation underlying memory (**Fig. 2C**). Confirming the smRNA-Seq results by qPCR in an independent cohort that was not sequenced and did not undergo extinction training, we found BLC mir-135b-5p levels were elevated in SEFL animals that displayed the greatest freezing one month earlier, during training (**Fig. 2D**). Further, the increase in mir-135b-5p was specific to the BLC, as no change was observed in other brain regions assessed from SEFL animals (**Fig. 2E**).

**Fig. 2.**
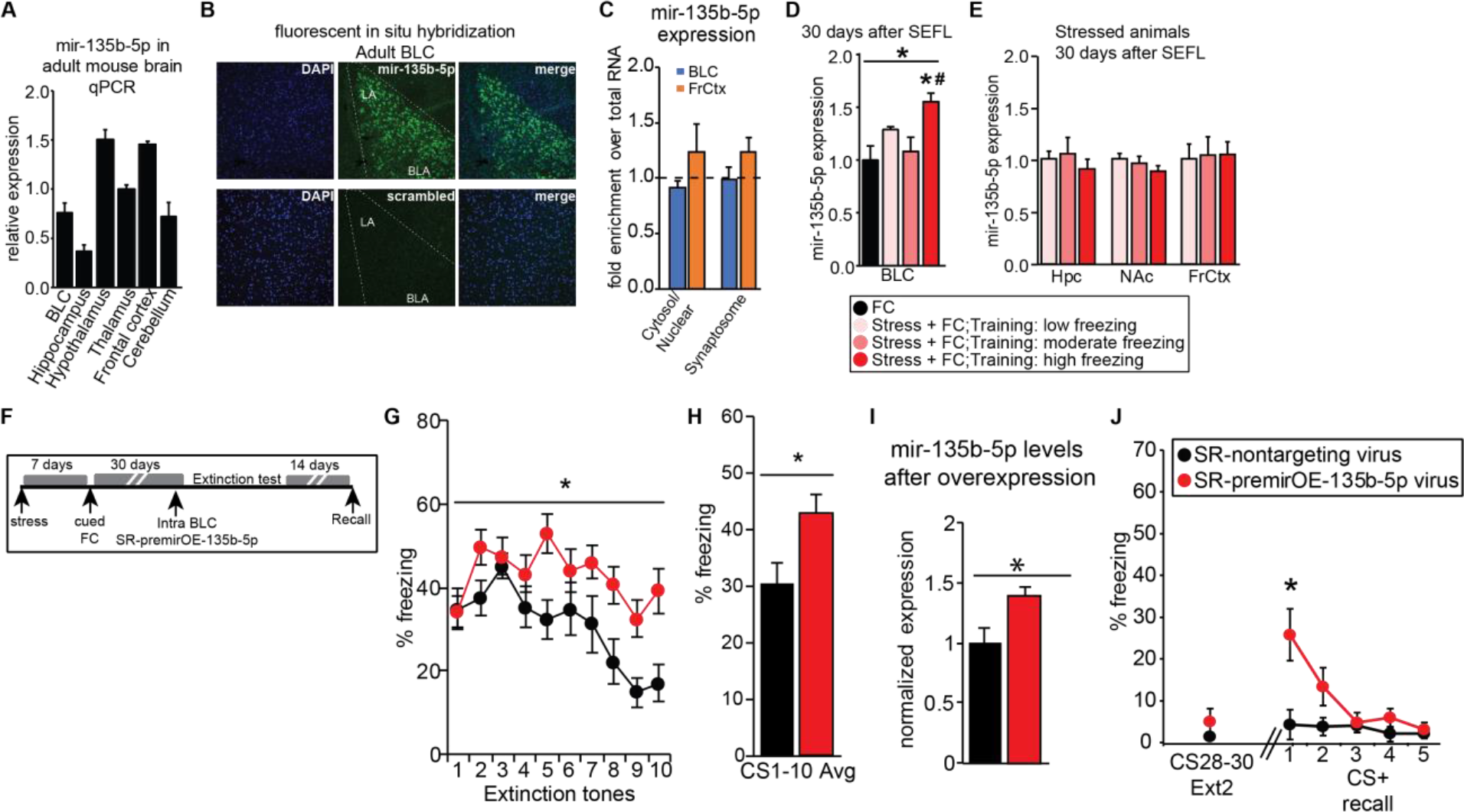
BLC overexpression of mir-135b-5p strengthens traumatic memory expression in SEFL mice. (A) mir-135b-5p expression in the adult mouse brain. N=3/region. (B) Localization of mir-135b-5p in the BLC measured with in situ hybridization. (C) mir-135b-5p expression in subcellular fractions to determine localization. N=4/region. (D) qPCR validation of elevated BLC mir-135b-5p expression in SS mice. One-way ANOVA: F_(3, 12)_ = 5.532; p=0.0128; Post-hoc Tukey test p<.05 for high vs FC (*) and high vs moderate (#). N=4-7/group. (E) mir-135b-5p levels in the hippocampus (Hpc), nucleus accumbens (NAc) or the frontal cortex (FrCtx) of mice 30 days after SEFL (no extinction). N=8-12/group. (F) Overview of strategy to OE BLC mir-135b-5p in SR animals. (G-H) Freezing during the first ten tones of extinction on Day 1. RM-ANOVA for 10 tones extinction day 1: F_(1,21)_=6.04; p=0.023. Unpaired t-test for average of 10 tones: t_(21)_ =2.457, p=0.0228. Nontargeting virus N=10; premirOE-135b-5p N=13. (I) mir-135b-5p levels in SR mice after extinction. Unpaired t-test: t_(18)_ =2.532, p=0.0209. Nontargeting virus N=9; premirOE-135b-5p N=11. (J) Extinction retention from the end of extinction day 2 (average freezing to tones 28-30) to a 5 tone recall test 2 weeks later. RM-ANOVA 5 tone recall: F_(1,20)_=5.27; p=0.033. Post-hoc t-test for tone 1: t_(20)_=2.673, p=0.0146. Nontargeting virus N=9; premirOE-135b-5p N=13.

We next examined the functional role of mir-135b-5p in regulating persistent stress-enhanced memory using premirOE-mir-135b-5p or a scrambled non-targeting sequence, as well as an anti-miRNA (sponge) viral construct that binds mir-135b-5p and prevents target binding. Injection of viruses into the BLC of naïve animals was localized, even at infusion volumes higher than those used for behavioral assays (**Fig. S3A**). We measured mir-135b-5p in the BLC 7 days after injection of OE or anti-miR constructs and, unexpectedly, observed increased mir-135b-5p in both conditions, without off-target effects on the levels of mir-135a-5p (**Fig. S3B**). miRNA expression is under tight homeostatic regulation and elevated mir-135b-5p after introduction of the sponge likely reflects compensatory upregulation to counter sequestration of mir-135b-5p. To mimic the elevated mir-135b-5p seen in SS mice, we injected premirOE-135b-5p into the BLC of SR mice 3 weeks after SEFL and performed extinction one week later (**Fig. 2F**). Enhanced memory expression was observed in SR-premirOE-135b-5p mice (**Fig. 2G-H**). Extinction was achieved with a strong protocol (**Fig. S4**), yet mir-135b-5p levels remained elevated at the conclusion of extinction in SR-premirOE-135b-5p mice relative to SR-nontargeting control animals (**Fig. 2I**), suggesting the potential for spontaneous recovery if mediated by mir-135b-5p. Indeed, a 5-tone recall test two weeks after extinction demonstrated a return of enhanced fear memory in SR-premirOE-135b-5p animals (**Fig. 2J**). Anxiety-related open field and eleveated plus maze behaviors were no different in naïve animals injected with premirOE-135b-5p in the BLC, indicating mir-135b-5p is not simply anxiogenic (**Fig. S5**).

The intent was to utilize the anti-mir-135b-5p virus to assess the impact of mir-135b-5p knockdown on stress-enhanced memory in SS mice. However, the compensatory response of mir-135b-5p to viral-mediated sequestration mimicked the premirOE-135b-5p virus (**Fig. S3**), elevating BLC mir-135b-5p levels. Consistent with this, intra-BLC anti-mir-135b-5p further exacerbated stress memory enhancement and produced hyperarousal (**Fig. S6**). Therefore, we instead used a synthetic hairpin mir-135b-5p inhibitor (INH) that sequesters mir-135b-5p, injecting it into the BLC of mice 28 days after SEFL, followed by extinction 2 days later (**Fig. 3A**). INH-mir-135b-5p reduced freezing in SS mice relative to a nontargeting control inhibitor during the first 10 CS+ tone presentations (**Fig. 3B-C**) and effectively inhibited mir-135b-5p expression (**Fig. 3D**). Following complete extinction (**Fig. S7**), we assessed extinction retention two weeks later in a 5-tone recall test and observed no spontaneous recovery in either group (**Fig. 3E**). However, a subthreshold training protocol resulted in a recovery of fear in SS-INH-nontargeting controls that was absent in SS-INH-mir-135b-5p mice, suggesting persistent dampening of memory via prior mir-135b-5p inhibition, rather than acute suppression of retrieval (**Fig. 3F**). Increased freezing from Recall Test 1 to 2 in SS-INH-nontargeting controls after subthreshold training was prevented in SS-INH-mir-135b-5p animals (**Fig. 3F**). BLC mir-135b-5p levels are not merely anxiotropic, as we observed no differences in anxiety-related tasks after INH-mir-135b-5p infusion in naïve mice (**Fig. S8**). Importantly, delivery of the nontargeting viral and inhibitor controls to the BLC did not disrupt expected phenotypic differences in memory strength between SS and SR animals (*4*) (**Fig. S9**), indicating that behavioral differences observed also cannot be attributed to technical manipulations of the BLC.

**Fig. 3.**
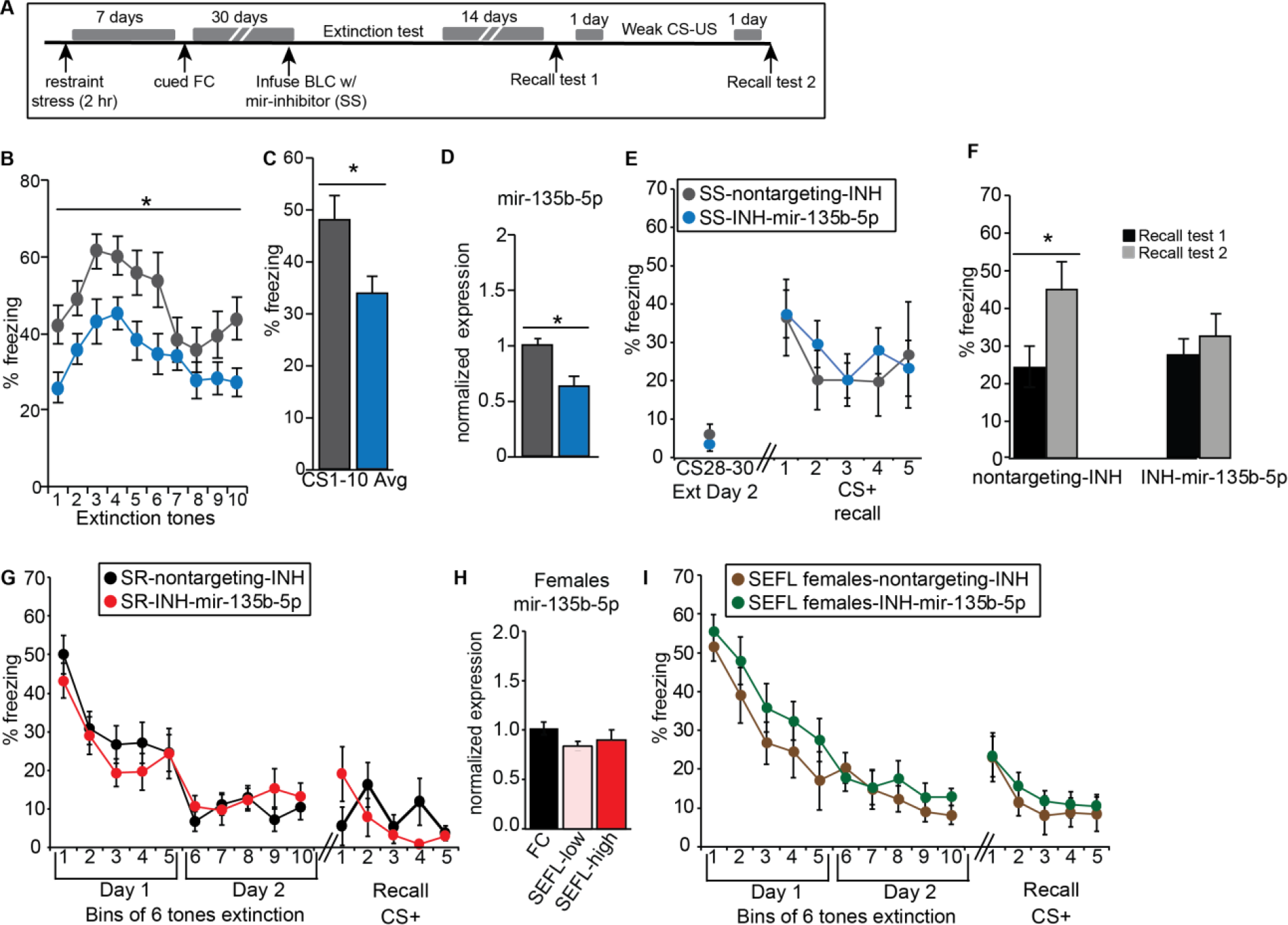
BLC inhibition of mir-135b-5p weakens stress-enhanced memory expression. (A) Overview of strategy to decrease BLC mir-135b-5p tone in SS animals. (B-C) Freezing during first 10 tones on Day 1 of extinction. RM-ANOVA for 10 tones extinction day 1: F_(1,28)_=5.814; p=0.023. Unpaired t-test for average of 10 tones: t_(29)_ =2.638, p=0.0133. Nontargeting control N=14; INH-mir-135b-5p N=17. (D) mir-135b-5p levels in SS mice after inhibition. Unpaired t-test: t_(18)_ =2.987, p=0.0079. Nontargeting control N=9; INH-mir-135b-5p N=11. (E) Extinction retention from the end of extinction Day 2 to the first 5 tone recall test. (F) Freezing after subthreshold re-training. Paired t-test SS-nontargetingINH: t_(3)_ =3.927, p=0.0294. Nontargeting control N=4; INH-mir-135b-5p N=6. (G) Extinction (6 tone bins) for SR male mice injected with INH-mir-135b-5p or nontargeting control. (H) mir-135b-5p expression in the BLC of SEFL females (no extinction) 30 days after training. (I) Extinction (6 tone bins) for stressed female mice injected with INH-mir-135b-5p inhibitor or nontargeting control.

Interestingly, further knockdown of BLC mir-135b-5p in SR animals via INH-mir-135b-5p infusion did not alter memory (**Fig. 3G**), as levels of mir-135b-5p are not different between FC and SR males (**Fig. 2D**). Consistent with our prior report (*4*), SEFL females do not cluster into stress susceptibility subgroups and levels of mir-135b-5p do not differ between FC and SEFL females, regardless of their freezing during training (**Fig. 3H**). Consistent with this, INH-mir-135b-5p in SEFL females does not reduce memory strength (**Fig. 3I**). Together this suggests a threshold of mir-135b-5p expression may have to be reached to achieve pathologic memory effects and subsequent amelioration by reduction of mir-135b-5p levels.

The mir-135b stem loop sequence is 100% conserved from mouse to human (**Fig. 4A**). While amygdala tissue from human PTSD subjects could not be obtained for this project, we measured levels of mature mir-135b-5p and the -3p arm, indicated to be the passenger strand by miRBase 22, in postmortem human amygdala by qPCR (**Fig. 4B and S10**). Both were detected, but consistent with miRBase 22, -5p expression was much higher levels than -3p, indicating -5p is the major mir-135b isoform in human amygdala. Using smRNA-Seq, we next measured levels of mir-135b-5p and -3p in serum samples collected from a cohort of well-characterized male Dutch military six months after a four month deployment in Afghanistan, at which time trauma exposure and PTSD symptomatology were assessed. mir-135b-3p was selectively elevated in members of the military diagnosed with PTSD (“susceptible”) relative to members without the diagnosis (“resilient”) and non-trauma exposed healthy military controls (**Fig. 4C-D and SE.4**). Passenger strands have traditionally been thought to be degraded upon release from the mature strand at the site of production. However, growing evidence indicates they can enter circulation and functionally impact distal structures (*29*). It will be important to identify the sites and actions of elevated mir-135b-3p. Regardless, this indicates mir-135b-3p’s potential as a biomarker of PTSD.

**Fig. 4.**
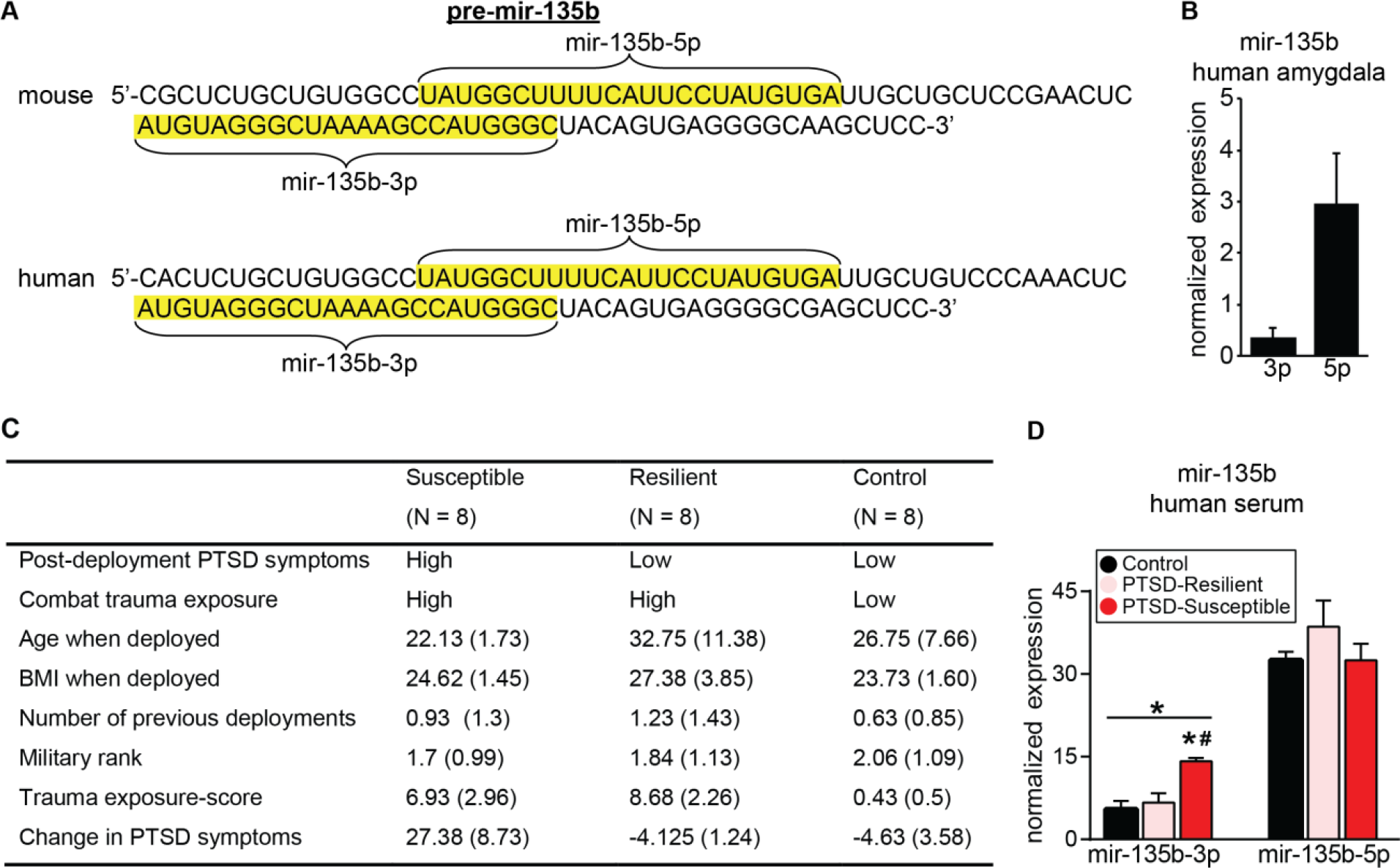
mir-135b is highly conserved in humans and elevated in the blood of PTSD-susceptible subjects. (A) Conservation of the mouse pre-mir-135b sequence from mouse to human. The sequences of the 5’ and 3’ arms are highlighted in yellow. (B) qPCR expression of mir-135b isoforms in human amygdala. N=4 brains. (C) Characteristics of human subjects used for the evaluation of circulating mir-135b in serum. (D) Sequencing expression of mir-135b isoforms in the serum of human subjects. One-way ANOVA for mir-135b-3p: F_(2, 17)_ = 15.32; p=0.0002; Post-hoc Tukey test p<.05 for control vs susceptible (*) and resilient vs susceptible (#). N=6-8/group.

Combining smRNA-Seq with quantitative MS identified putative miRNA-dependent and -independent pathways selectively dysregulated in SS animals. Integration of such datasets is unprecedented in the memory field and led to identification of mir-135b-5p as a critical regulator of remote, stress-enhanced memory strength. This work may be therapeutically relevant to individuals that experience long-lasting traumatic memories and sheds light on the potential of miRNAs to critically regulate essential neurological processes, such as memory persistence.

Characterization of mir-135b-5p in the brain demonstrated that it is poised to regulate stress-related memory in amygdala neurons. The immediate behavioral effects of mir-135b-5p overexpression in SR and knockdown in SS mice at the onset of memory retrieval and the overall normal extinction rate over two days indicate it is unlikely that mir-135b-5p inhibition disrupts memory through accelerated extinction. Alternatively, mir-135b-5p could have an acute effect on retrieval of remote fear memory. However, inhibition of mir-135b-5p in SS animals prevented the spontaneous recovery of fear after subthreshold training an entire month after administration of the inhibitor, suggesting that mir-135b-5p may selectively participate in storage of stress-enhanced memories.

Development of a small molecule inhibitor of the mir-135b precursor (*30*) would provide the temporal resolution necessary to assess its role in memory storage. But more importantly, it would have potential therapeutic value. The animal studies presented here indicate that mir-135b-5p inhibition is highly selective, only impacting memory when the miRNA is elevated (**Fig. 3**). Further, the increase in circulating levels of mir-135b-3p specific to members of the military with a PTSD diagnosis suggests it may serve as an important biomarker capable of identifying individuals that would be responsive to a mir-135b-based therapeutic.

## Acknowledgements

We thank the Scripps Florida Genomics Core for sequencing services, Nripesh Prasad at the Genomic Services Lab at Hudson Alpha for sequencing services and data analysis, Adrian Reich and the Bioinformatics Core for data analysis, the Mouse Behavior core and Alicia Brantley for assistance and behavioral equipment, all members of the Miller/Rumbaugh Labs for their technical assistance and thoughtful discussions.

## Author contributions

SES designed and performed experiments, analyzed data and wrote the manuscript; SJ, MJ, CS, TK, and NFJ performed experiments and analyzed data; JK and JK analyzed data; LdN, CHV, MPMB, EG and EV designed and performed experiments, analyzed data; KJR and GR wrote the manuscript; BPFR designed experiments, analyzed data; and CAM designed the study, analyzed data and wrote the manuscript.

